# α-Synuclein aggregates in corticostriatal terminals impair glutamatergic transmission in the absence of neurodegeneration

**DOI:** 10.64898/2026.08.03.742532

**Authors:** Charlotte F. Brzozowski, Zoe N. Fokakis, Marissa A. Menard, Harshita V. Challa, Ignacio Gallardo, J. Dominic Hall, Douglas Narbert, Michael Millet, J. Andrew Hardaway, Mark S. Moehle, Laura A. Volpicelli-Daley

**Affiliations:** Department of Neurology, Killion Center for Neurodegeneration and Experimental Therapeutics, University of Alabama at Birmingham, Birmingham, AL, 35294, USA; Department of Psychiatry and Behavioral Neurobiology, University of Alabama at Birmingham, Birmingham, AL, 35294, USA; Department of Pharmacology and Therapeutics and Center for Translational Research in Neurodegeneration, University of Florida, Gainesville, FL, 32610, USA; Aligning Science Across Parkinson’s (ASAP) Collaborative Research Network, Chevy Chase, MD, 20815

## Abstract

Substantia nigra pars compacta dopamine neuron loss and Lewy pathology, aggregates of α-synuclein, characterize Parkinson’s disease and Dementia with Lewy Bodies. Lewy pathology localizes to cortical neurons, and is found as Lewy neurites in the striatum, but its effects on excitatory synaptic function are just beginning to be understood. Corticostriatal projections regulate motor and cognitive behaviors impaired in these disorders. Here, α-synuclein aggregation was induced in mouse M2 cortex, a vulnerable region in human disease. Early after initiation, aggregates localized to corticostriatal vesicular glutamate transporter 1 (vGLUT1)-positive terminals, with sparing of spiny projection neuron (SPN) soma, and dopamine terminals and neurons. Corticostriatal presynaptic aggregates significantly impaired glutamatergic transmission, without overt cortical neuron loss, and were associated with decreased synaptic density and volume. Thus, formation of presynaptic α-synuclein aggregates impairs corticostriatal function without degeneration of cortical neurons or striatal dopamine terminals, suggesting pathologic α-synuclein is sufficient for synaptic loss. Our findings also point to early synaptic dysfunction as a therapeutic target in Lewy body diseases.

## Introduction

In Parkinson’s disease (PD) and Dementia with Lewy bodies (DLB), degeneration of the nigrostriatal pathway causes motor symptoms which are alleviated by dopamine replacement therapies such as L-DOPA. Cognitive impairments also occur and can be mitigated by cholinesterase inhibitors. However, neither L-DOPA nor cholinesterase inhibitors prevent the progression of Lewy body diseases (LBD), and many motor and non-motor symptoms remain refractory to treatment. Thus, it is important to understand disease processes in additional brain circuits as well as the mechanisms leading to neuronal dysfunction. Lewy pathology composed of pathologic aggregates of α-synuclein (α-syn) localizes to specific brain regions where it correlates with LBD symptoms. Specifically, cortical neurons show widespread α-syn pathology.^1,2^ Projections from the cortex to striatum regulate motor and cognitive behaviors and have been implicated in PD motor and non-motor symptoms. Glutamatergic projections from the cortex synapse on striatal spiny projection neurons (SPNs). Both in post-mortem pathologic studies of PD striatum and rodent dopamine depletion models of PD, there is a decrease of dendritic SPN spines, reflecting a loss of striatal glutamatergic synapses.^3,4^

The pre-supplementary motor area (pre-SMA; M2 in mice) shows vulnerability in LBDs. Imaging studies of PD patients show reduced activity of the pre-supplementary motor cortical area during movement.^5^ Pathological studies show that the pre-SMA is the only cortical area with neuron loss in PD.^6^ The pre-SMA sends dense projections to the striatum, and transmagnetic stimulation, a non-invasive method to target cortical areas, of the pre-SMA improves PD motor symptoms.^7,8^ In addition to movement, the pre-SMA integrates sensory information and motivation in the completion of complex, goal-directed behaviors, which are impaired in PD.^9–13^ Therefore, studying pre-SMA/M2 to striatal connections and potential impairment in LBDs could help us understand how to improve both motor and non-motor symptoms.

In the striatum, Lewy neurites, axonal accumulations of α-syn aggregates, are the primary form of Lewy pathology.^14^ How Lewy pathology impacts corticostriatal transmission has only begun to be studied. Injections of α-syn pre-formed fibrils (PFFs) into the mouse striatum induces endogenous α-syn to aggregate and allows studies of the consequences of aggregate formation on neuron function. At an early six-week time point after intrastriatal injections of PFFs, electrophysiological recordings at SPNs in the striatum show reduced corticostriatal glutamate drive, intrinsic excitability, and asynchronous release frequency suggesting reductions in presynaptic release sites.^15^ Aggregates of α-syn appeared in vGLUT1-positive corticostriatal terminals and in SPN soma, as well as dopamine neurons of the substantia nigra pars compacta. In addition, dopamine terminals were significantly reduced six weeks following striatal PFF injections compared to controls. Additional studies have implicated that exposure of striatum to α-syn oligomers alters striatal plasticity in a dopamine dependent manner.^16^ Therefore, the corticostriatal synaptic impairments observed could be caused by multiple mechanisms including aggregate specific effects within cortical neurons, aggregates in somatodendritic domains of SPNs, or reduced dopamine terminals. This then makes understanding the consequences of α-syn aggregation specifically in excitatory corticostriatal terminals difficult.

In contrast to striatal PFF injections, injections of PFFs into the cortex produce aggregates in interconnected cortical brain regions but not in the substantia nigra pars compacta.^17^ Aggregates can spread from the soma anterogradely to terminals.^18,19^ Cortical injections could then produce α-syn pathology in cortical projection neuron terminals without confounding effects of pathology in SPNs or in dopamine neurons. Here, injections of PFFs into M2 cortex produced α-syn aggregates in vGLUT1-positive corticostriatal terminals without SPN or substantia nigra pars compacta pathology. Thus, the effect of presynaptic abnormal α-syn on synaptic function could be determined without somatodendric SPN pathology as well as without loss of nigrostriatal dopamine, allowing us to mechanistically dissect how Lewy pathology specifically alters M2 function. Within this cortical predominant Lewy pathology model, we utilized a multi-disciplinary approach to rigorously dissect the effects of α-syn aggregation on cortical glutamatergic terminals in the striatum. To determine the impact of α-syn on synaptic physiology, we utilized an optogenetic approach using ChrimsonR expressed in M2 cortex to specifically examine synaptic efficacy of M2 cortical synapses in the striatum. We found stimulation of M2 corticostriatal terminals produced robust excitatory postsynaptic potentials in control mice but not in PFF injected mice and did not change SPN intrinsic excitability. Three-dimensional reconstructions of corticostriatal synaptic loci revealed that reduced synaptic density and volumes at least partially account for reduced evoked glutamate release, which was confirmed through *ex vivo* electrophysiological asynchronous release studies. Thus, at early stages after formation, presynaptic α-syn pathology is sufficient to cause a robust impairment of corticostriatal function which is intrinsic to the formation of pathology in these neurons and is not influenced by decreased dopamine terminals or pathology formation in SPNs.

## Results

### M2-PFF injections cause p-**α**-syn-positive aggregate accumulation in cortex and amygdala but not substantia nigra pars compacta

The goal of this project was to determine the effect of PFF injections in M2 cortex on corticostriatal excitatory synapses at an early time point, six weeks, after injections without the confounding effects of changes in dopamine neurons or pathology in post-synaptic neurons. PFFs were injected directly into M2 cortex unilaterally. Analyses of pathologic α-syn aggregates were performed six weeks after injections, when α-syn aggregates begin to appear in neurons in the brain after PFF exposure. α-Syn pathology was visualized using an antibody directed at phosphorylated α-syn at serine 129 (p-α-syn).^20^ Immunofluorescence showed α-syn aggregates both ipsilateral and contralateral in secondary motor cortex (M2), orbital area (ORB), and primary motor cortex (M1) (Figure 1a,b). Aggregates also formed in the basolateral amygdala (BLA), consistent with connections from BLA and M2.^21^ Aggregates were not obvious in the dorsal striatum at low magnification (Figure 1a) but were visible at higher magnification (Figure 1b) and resembled Lewy neurite threads or small puncta. Aggregates of p-α-syn did not form in the substantia nigra pars compacta at six weeks post-injection. Monomeric α-syn control injections into M2 did not cause pathologic p-α-syn aggregation (Supplemental Figure 1).

**Figure 1:**
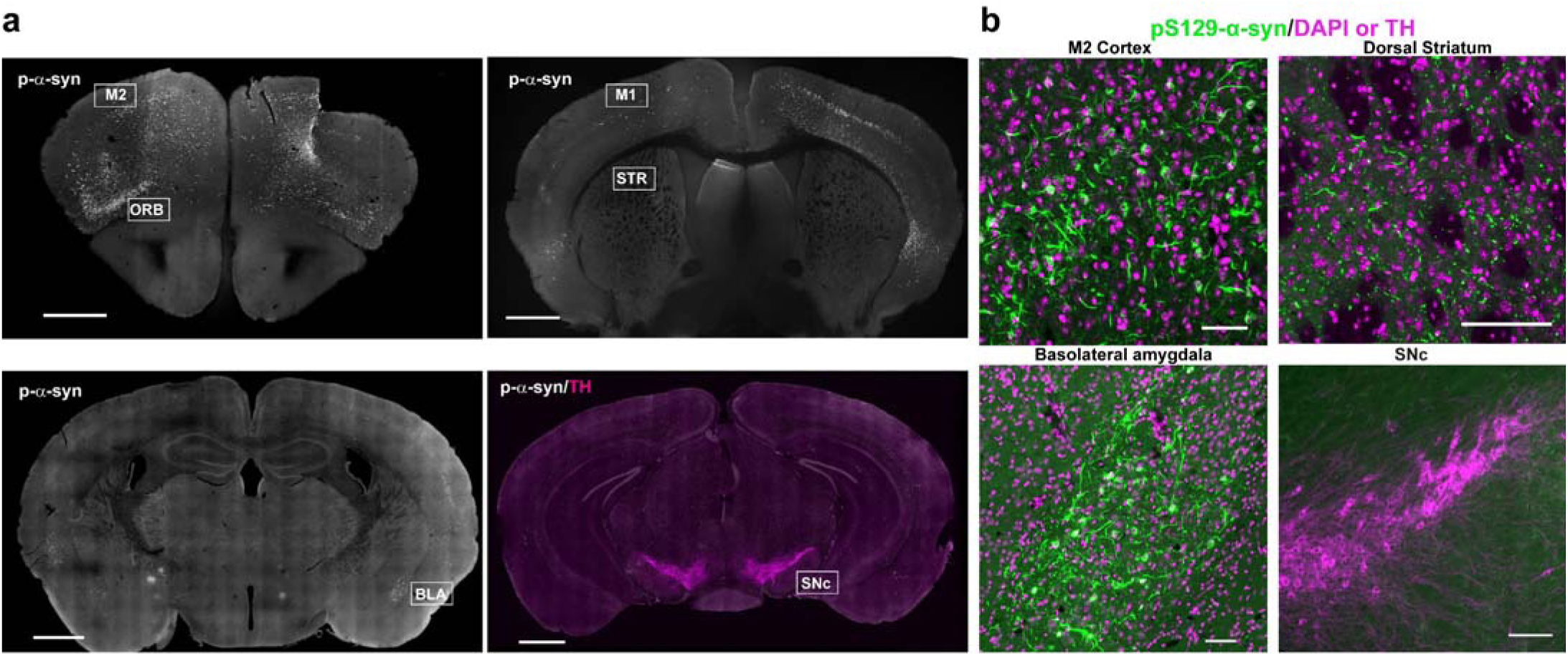
M2-PFF injection causes cortical-dominant p-α-syn pathology. Mice were injected unilaterally with 10 µg PFFs in M2 cortex and were perfused six weeks later. Immunofluorescence using an antibody for p-α-syn was performed on 40 µm coronal sections to identify α-syn aggregates. **a)** Representative confocal images showing localization of p-α-syn (white). The bottom right section shows tyrosine hydroxylase immunofluorescence (magenta) to visualize the substantia nigra pars compacta. Scale bars: 1,000 µm. The ipsilateral injection site is on the right side in each panel. Abbreviations: M2 = Secondary Motor Cortex, ORB = Orbital Area, M1 = Primary Motor Cortex, STR = striatum, BLA = Amygdala, SNc = substantia nigra pars compacta. **b)** Fluorescent images of p-α-syn pathology (green) and either DAPI (magenta) to visualize nuclei or TH (magenta, bottom right) to visualize the substantia nigra pars compacta of selected brain areas depicting higher magnification from panels a, Scale bars: 100 µm.

### Accumulation of p-**α**-syn pathology in cortical layer V neurons does not cause early neuron loss

Previously, it was shown that PFF injections into the striatum lead to accumulation of p-α-syn aggregates in cortical layer V projection neurons.^22,23^ This pattern of pathology was preserved with M2 injections; p-α-syn inclusions localized predominantly to cortical layer V pyramidal neurons (Figure 2a). Aggregates were visible ipsilateral and contralateral to the injection site, in line with known inputs into M2. This pattern was visible in M2 and other cortical areas such as primary motor cortex, primary sensory cortex, and the auditory cortex (Supplemental Figure 2).

**Figure 2:**
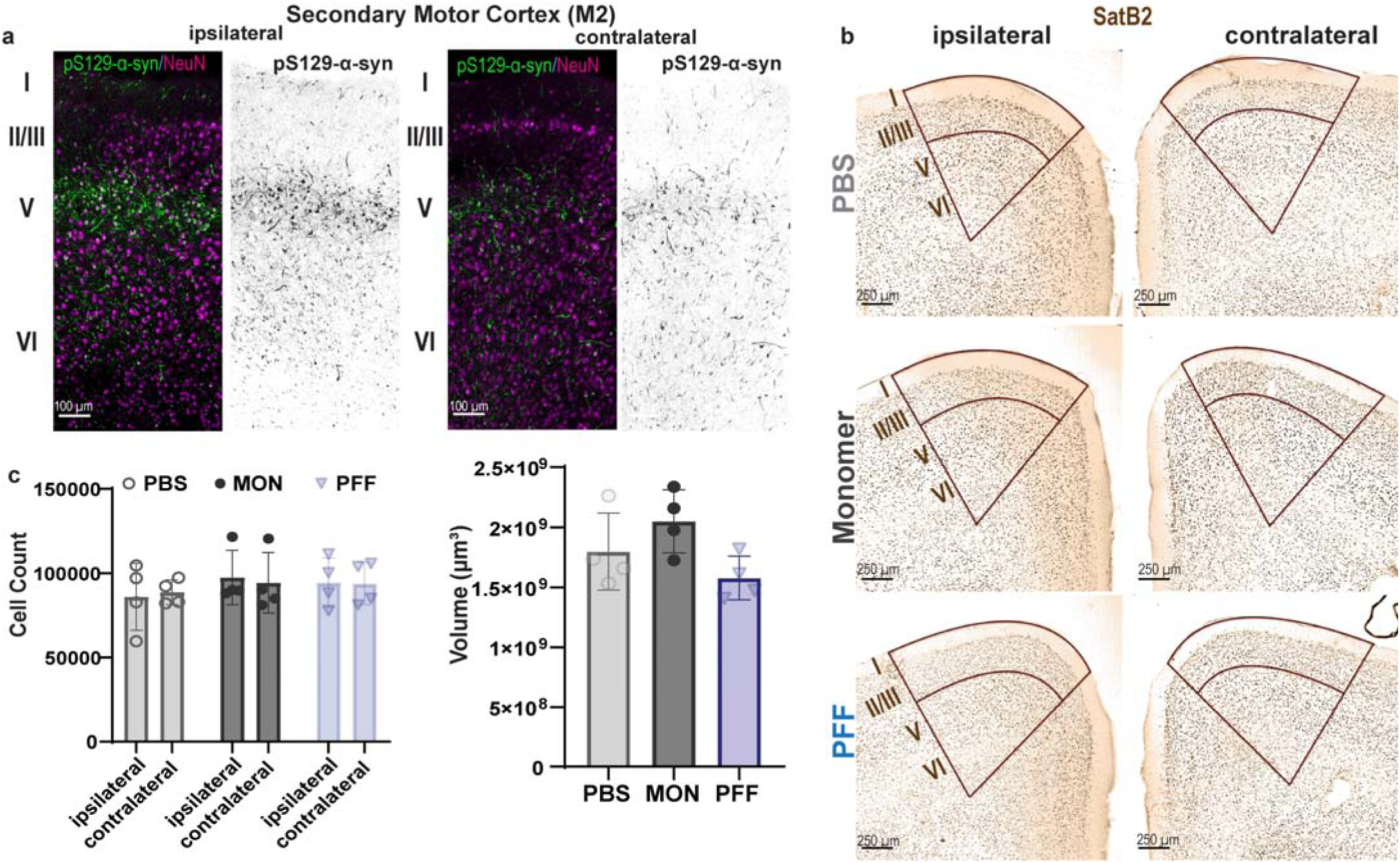
Cortical p-α-syn pathology accumulates in layer V neurons. Mice were injected with 10 µg PFFs, or monomer, or PBS in M2 cortex and were perfused six weeks later. **a)** Shown are confocal images of p-α-syn aggregates (green) and the neuronal marker NeuN (magenta) in M2 cortex ipsilateral (left two panels) and contralateral (right two panels) to the M2 injection site. Inverted black and white images are shown next to the fluorescent images to better visualize the p-α-syn aggregates. Scale bars = 100 µm. **b)** Shown are representative brightfield images of coronal sections stained for SatB2 for unilateral PBS, monomer, and PFF injected mice. The area outlined in brown represents the outline of the M2 of ipsilateral (left) and contralateral side (right), and the different cortical layers are annotated on the ipsilateral side. Deep layer V+VI are further outlined. Scale bars=250 µm. **c)** Quantification of the deep layer (V+VI) SatB2 neuron cell count using unbiased stereology showed no significant changes between injected and non-injected hemispheres, nor between groups. (Two-way ANOVA: F(2,18)=0.07, p=0.93). Volume of outlined secondary motor cortices for stereology did not show differences among groups (One-way ANOVA, F(2, 9) = 3.281, p=0.0851).

To assess whether M2 injections of PFFs and subsequent cortical α-syn pathology formation cause neuron loss in the cortex, immunohistochemistry was performed using an antibody to the excitatory neuronal marker, SatB2.^22^ Unbiased stereology was performed to quantify SatB2-positive neurons in M2 cortex from mice injected with PFFs or phosphate buffered saline (PBS), or monomeric α-syn as controls. Quantitation of cortical layers V and VI in M2 showed no differences in neuron count among treatment groups at six weeks following PFF injections into M2 (Figure 2b, 2c), as has been previously reported for cortical neuron counts with striatal injections of PFFs.^22^

### Striatal pathology is not associated with dopamine terminal loss and localizes to corticostriatal terminals

Analyses of dop amine transporter (DAT) immunofluorescence in the dorsal striatum of mice with M2 injections of PBS, monomer, or PFFs showed no loss of dopamine terminals six weeks post-injections, unlike PFF injections into the striatum (Figure 3a,b).^15^ In mice injected with PFFs into M2 cortex, striatal p-α-syn pathology was neuritic (Figure 3c). Double labeling immunofluorescence of p-α-syn and the SPN marker DARPP-32 showed no somal p-α-syn aggregates in SPN cell bodies (Supplemental Figure 3) in M2-PFF injected mice, unlike PFF injections into the striatum which show robust somal SPN pathology.^15^ Neuritic p-α-syn pathology in the striatum overlapped with vGLUT1 in corticostriatal terminals (Figure 3c). Only a small proportion of p-α-syn overlapped with DAT at nigral-striatal dopamine terminals (Figure 3d). Colocalization analyses of p-α-syn and vGLUT1 compared to p-α-syn and DAT showed significantly more overlap between p-α-syn pathology and vGLUT1 than overlap with DAT-positive terminals (Figure 3e).

**Figure 3:**
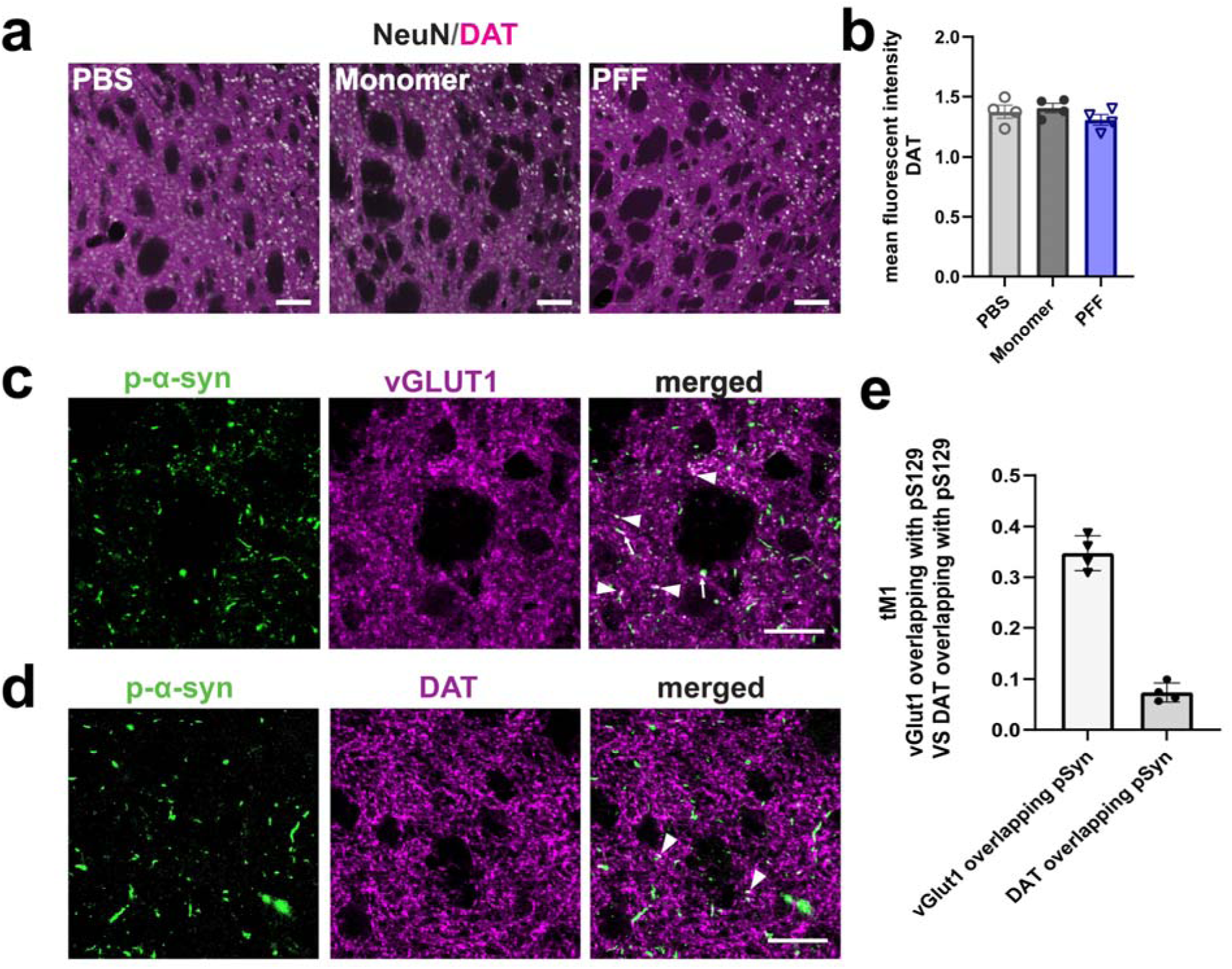
Neuritic, striatal p-α-syn pathology overlaps with the corticostriatal, presynaptic marker, vGLUT1. Mice were injected with 10 µg PBS, monomer, or PFFs in M2 cortex and were perfused six weeks later. **a)** Confocal images of dorsal striatum immunofluorescence for DAT (magenta) and NeuN (white). Scale bar = 100 µm. **b)** Fluorescence intensity of DAT in the dorsal medial striatum ipsilateral to the M2-PFF injection site was normalized to fluorescence of NeuN. One-way ANOVA was not significant F(2,9) =1.18, p=0.35. **c)** Dorsal striatum immunofluorescence for p-α-syn pathology (green) and synaptic marker vGLUT1 (magenta). Arrowheads point to colocalization (white). Scale bar = 100 µm. **d)** Dorsal striatum immunofluorescence for p-α-syn pathology (green) and DAT (magenta). Arrowheads point to colocalization (white). Scale bar = 100 µm **d)** Quantification of colocalization analyses using thresholded Manders (tM1) Colocalization Coefficient between p-α-syn pathology and glutamatergic terminal marker vGLUT1 and between p-α-syn pathology and DAT. Independent t-test t=15.18, p<0.0001.

### Impaired M2-to-striatum glutamate transmission in mice with corticostriatal **α**-syn aggregates is driven by a reduction in active corticostriatal release sites

To determine the impact of M2 specific pathology on cortical to SPN synapse physiology, AAV9-hSyn-ChrimsonR-tdTOMATO, expressing the red-shifted channelrhodopsin ChrimsonR was co-injected into M2 cortex along with PFFs or monomeric α-syn. To assess the optogenetically-evoked glutamate transmission of M2-to-striatum projections, whole cell patch clamp recordings of SPNs in the ipsilateral dorsal striatum within regions positive for tdTomato expression were performed six weeks after M2-PFF or monomer injections (Figure 4a). Using a 617 nm LED light (ThorLabs) to evoke glutamate transmission onto SPNs, the optogenetically evoked excitatory postsynaptic currents (oeEPSC) in response to increasing optogenetic stimulation intensities were measured. Monomer control injected mice showed increased oeEPSCs with increased stimulation intensity (Figure 4b,c). However, M2-PFF injected mice showed significantly reduced oeEPSC amplitudes compared to control mice. Using a paired-pulse stimulation paradigm of optical stimuli (50 ms interval) to assess pre-synaptic release probability, we observed no change in paired-pulse ratio in M2-PFF injected mice (Figure 4d) compared to controls, suggesting no change in glutamate release probability. Recordings of total spontaneous EPSCs in SPNs showed no difference between PFF and monomer injected mice with respect to frequency or amplitude of total spontaneous excitatory transmission (Supplemental Figure 4). Asynchronous release experiments were performed to assess whether the observed decrease in oeEPSC amplitude in M2-PFF injected mice resulted from changes in the number of active release sites or changes to synaptic strength. Desynchronization of EPSC into asynchronous EPSCs (aEPSC) was facilitated by substituting Ca^2+^ with Sr^2+^ ions in the bath chamber and were optically evoked as above (oeaEPSC). Asynchronous release experiments for M2-to-striatum specific transmission revealed a significant increase in the inter event interval between oeaEPSCs (Figure 4e,f) in M2-PFF injected mice compared to M2-monomer injected mice, showing a reduced frequency of oeaEPSC, and suggesting a decrease in release sites. aEPSC amplitude was not changed between groups (Figure 4e,g).

**Figure 4:**
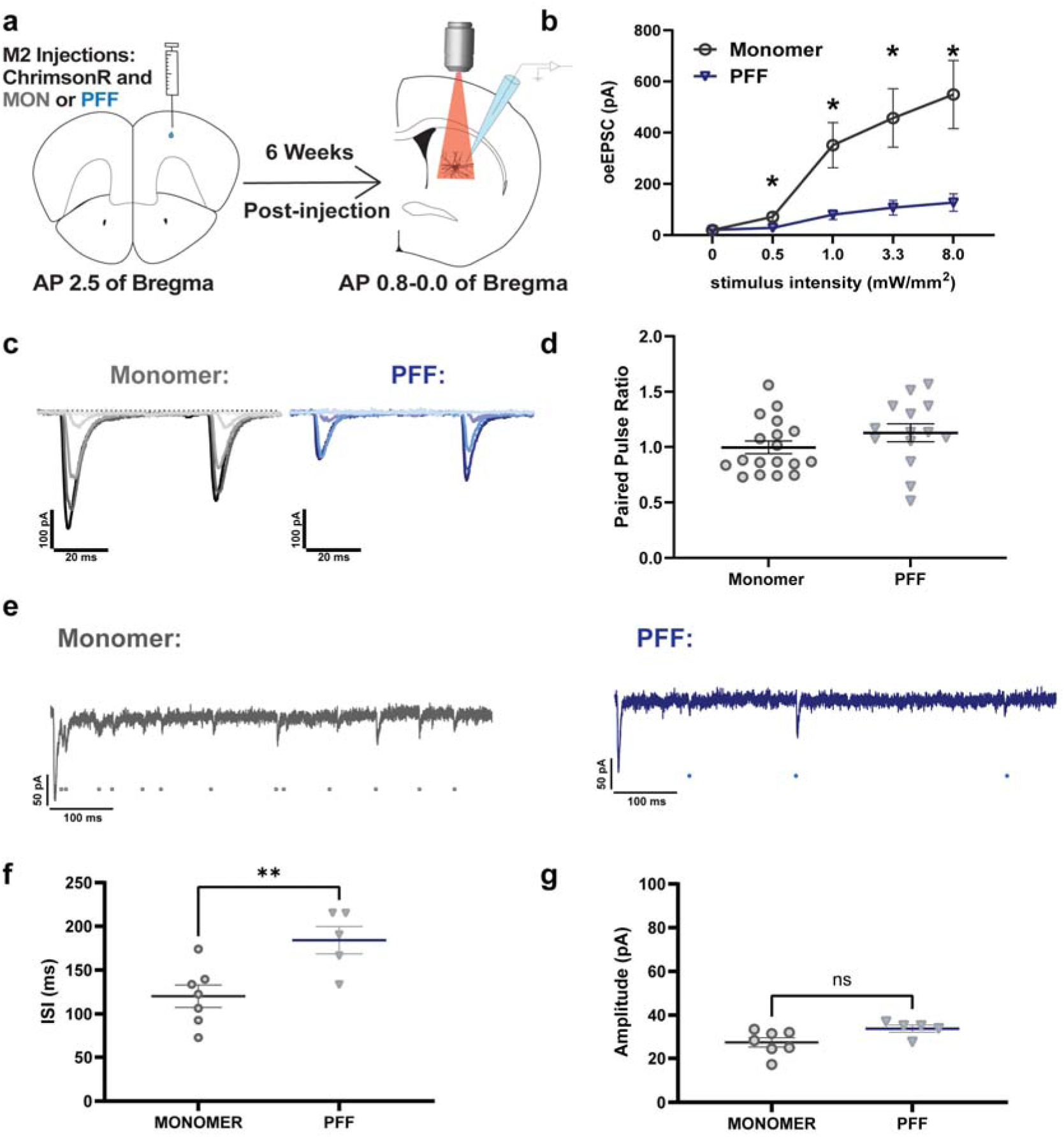
Impaired M2-terminal specific, optogenetically evoked glutamate transmission and decrease in active release sites onto SPNs in p-α-syn pathology positive mice. **a)** Schematic illustration (Created with BioRender.com) of experimental setup: 3-month-old C57BL/6J mice received unilateral M2 cortex injections of PFFs or monomeric LJ-syn as control, in addition to AAV9-hSyn-ChrimsonR-tdTomato. Whole-cell patch clamp recordings of SPNs were performed six weeks post-injections after identifying tdTomato positive terminals in the dorso-medial striatum ipsilateral to injection site. **b)** Input-output curves generated by stimulating ChrimsonR-positive M2 terminals with far red shifted light with increasing LED light intensity and measuring the optogenetically-evoked postsynaptic current in SPNs. Data did not fit a normal distribution and were transformed log_10_ which achieved normality. Data were analyzed with two-way repeated measures ANOVA with Geisser-Greenhouse correction. Monomer (grey): n=14, N=4; PFF (blue): n=12, N=4. Two-way Repeated Measure ANOVA analysis with Geisser-Greenhouse correction, Interaction: F(2, 40)=14.6; p<0.0001. Šidák’s multiple comparison post hoc was performed. Data presented are the non-transformed mean pA (±SEM) (expressed as positive values for ease of visualization). **c)** Representative oeEPSC traces for ChrimsonR optogenetic paired pulse stimulation with increasing stimulus intensity for monomer injected mice (gray) and PFF injected mice (blue); higher stimuli are indicated by darker colors. **d)** Paired pulse stimulation (50 ms interval) at 3.3 mW stimulation was calculated as the ratio of the pA of stimulus 2 divided by pA of stimulus 1. Monomer: n=18, N=4; PFF: n=14, N=4. Independent t test=1.3, p=0.20. **e)** Representative traces of optogenetically-evoked Sr^2+^ asynchronous release EPSC for monomer (gray) and PFF (blue) injections. Sr^2+^ oeEPSCs are indicated by circles below the representative traces. **f)** Quantification of the mean inter event interval (IEI) of asynchronous EPSCs for monomer and M2-PFF injected mice. Monomer: n=7, N=4; PFF: n=5, N=3. Independent t test=3.2, p=0.009. **g)** Quantification of the mean asynchronous EPSC amplitude (presented as positive pA) between monomer and M2-PFF injections. MON: n=7, N=4; PFF: n=5, N=3. Independent test=2.2, p=0.055.

To assess whether any of the observed changes might be due to changes of intrinsic excitability, the rheobase current in SPNs of the different treatment groups was measured. No changes in rheobase current or other measures of intrinsic excitability between groups (Supplemental Figure 5) were observed. Additionally, no changes in passive membrane properties of SPNs were observed between M2-PFF injected mice and control mice (Supplemental Figure5, unlike SPNs from striatal SPN injected mice.^15^

### Cortical pathology formation causes a reduction in corticostriatal synaptic loci density

Using a three-dimensional surface rendering approach developed by our lab previously,^24^ corticostriatal synaptic loci in the ipsilateral dorso-central striatum of six-weeks post unilaterally M2-PFF injected mice, or control injected mice (PBS or monomeric α-syn) were quantified. Corticostriatal synapses were visualized by immunofluorescence for the excitatory, cortical terminal marker vGLUT1 and the excitatory, postsynaptic marker Homer1. Using Imaris analysis software and its ‘surface’ function, 3D surfaces for vGLUT1 and Homer1 were generated based on fluorescent signal in confocal z-stacks. To filter for synaptic loci, the shortest distance (<0.01µm) filter between reconstructed vGLUT1 and Homer1 surfaces was used to isolate synaptic pairs. Representative images of confocal images, and Imaris 3D reconstructed synaptic loci are shown in Figure 5a. The density of both synaptic vGLUT1 and Homer1 positive synaptic loci in the ipsilateral dorso-medial striatum was significantly reduced six weeks after unilateral M2-PFF injections compared to controls, with no change in density observed between the PBS or monomer groups (Figure 5b). Synaptic Homer1 surface volumes were also significantly reduced in M2-PFF injected mice compared to monomer α-syn control mice (Figure 5b).

**Figure 5:**
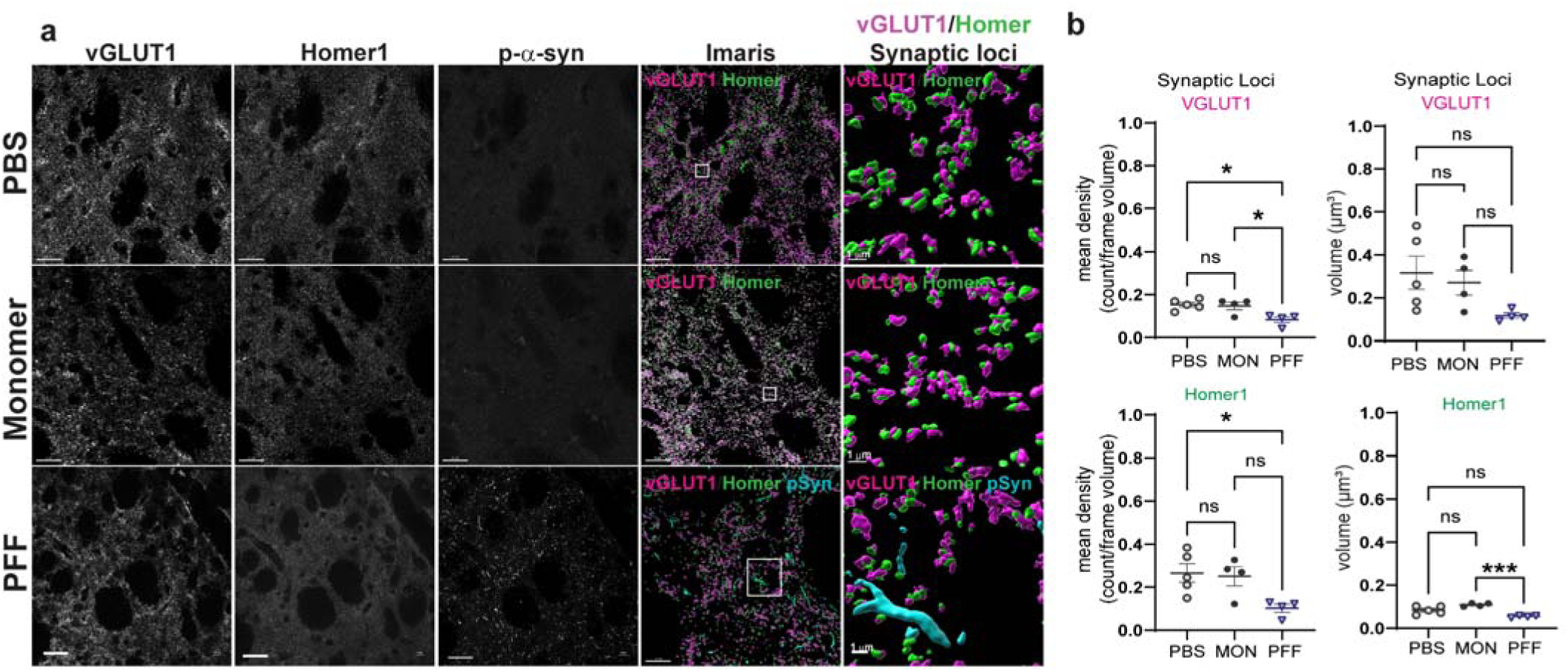
M2-PFF injections cause reduction in vGLUT1/Homer1 synaptic loci. Mice were injected with 10 µg PFFs or monomer or PBS in M2 cortex and were perfused six weeks later. **a)** Immunofluorescence was performed for the presynaptic corticostriatal marker vGLUT1, excitatory postsynaptic marker Homer1, and p-α-syn. All confocal images were captured with same laser intensity. Images were deconvolved. Imaris was used to perform 3D rendering and filtering of synaptic loci (vGLUT1 juxtaposed to Homer1 <0.01 µm). The first three columns are black and white confocal images of vGLUT1, Homer1, and p-α-syn, respectively. The fourth column shows the Imaris rendered, synaptic loci filtered image of vGLUT1 (magenta), Homer1 (green), and p-α-syn (cyan; bottom row, PFF). The fifth column of images shows Imaris rendered vGLUT1/Homer1 surfaces in close proximity. Scale bar on left = 1 or 20 µm.sarala**b)** Quantification of density of synaptic loci vGLUT1 and Homer1 surfaces. Synaptic loci in the 3D rendered image filtered for synaptic loci. The counts were divided by frame volume to calculate mean density. Volumes of synaptic vGLUT1 or Homer1 were determined by Imaris. Synaptic vGLUT1 density: One way ANOVA, nested design with frames nested within individual mice. F(2,10)=7.5, p=0.0103, Tukey’s posthoc test PBS vs. PFF p=0.0123, monomer vs. PFF p=0.027. Synaptic vGLUT1 volume F(2,10)=3.0, p=0.0955. Synaptic Homer1 density: One way ANOVA nested design with frames nested within individual mice. F(2,10)=5.3, p=0.0268, Tukey’s posthoc test PBS vs. PFF p=0.0309. Synaptic Homer1 volume F(2,10)=5.7, p=0.0221. Tukey’s posthoc test monomer vs. PFF p=0.0214.

## Discussion

The accumulation of intraneuronal Lewy pathology is a pathognomonic feature of synucleinopathies such as PD and DLB. Aberrant corticostriatal transmission could contribute to PD symptoms such as impaired initiation of movement, errors in serial order tasks, executive dysfunction, and impaired reinforcement learning.^11,25,26^ Dopamine depletion models of PD show that loss of striatal dopamine decreases excitatory drive onto spiny projection neurons, but these models do not account for pathologic α-syn, another hallmark of PD.^3,27,28^ In order to analyze the impact of abnormal α-syn, intrastriatal injections of α-syn oligomers or PFFs have been performed, and disrupt corticostriatal activity. However, dopamine terminal loss occurs early in the intrastriatal models,^15^ and thus reduced dopamine may still contribute to synaptic defects. In addition, in the intrastriatal PFF model, α-syn aggregates form in the soma of spiny projection neurons, making it difficult to determine if impaired synaptic transmission results from presynaptic or postsynaptic pathology. Since α-syn is a presynaptic protein, aggregation initiates in terminals and axons,^19,29^ and in PD, the most abundant form of Lewy pathology in the striatum is neuritic, suggesting that the majority of Lewy pathology in the human striatum is found in terminals arising from neurons projecting to the striatum.^14^ We therefore sought to restrict formation of α-syn aggregates to corticostriatal terminals by injecting PFFs directly into M2 cortex, a cortical area particularly susceptible in PD.^6^ We showed aggregated p-α-syn was restricted to vGLUT1-positive corticostriatal terminals but was not observed in soma of SPNs or in the substantia nigra pars compacta. Unlike the intrastriatal PFF model in which dopamine terminal immunofluorescence is reduced as early as six weeks after PFF injections,^15^ there was no significant loss of dopamine terminals six weeks following unilateral M2 injections of PFFs. We observed a pathological phenotype indicative of functional synapse loss, as evidenced by significantly reduced oeEPSCs and reduction in frequency of oeaEPSCs in *ex vivo* slice electrophysiological recordings. Thus, pathologic α-synuclein is sufficient for impaired corticostriatal transmission.

At the early timepoint after aggregate initiation analyzed in this study, no neuron loss is observed in deep layers V and VI of the cortex or throughout the M2 cortex. In addition, amplitude of aEPSCs was unchanged in PFF or MON injected animals, indicating quantal size of synaptic vesicles is likely unaffected. This is consistent with our previous work showing no changes in docked vesicles within excitatory synapses of the amygdala or cortex of PFF injected mice.^30,31^ Deficits in excitatory neuronal activity, therefore, are likely indicative of changes within the synapses at the presynaptic terminal of the corticostriatal glutamatergic projections. α-Syn induced alterations in corticostriatal synaptic function specifically have thus far only been explored in the context of dopaminergic loss.^16^ However, in this study synaptic dysfunction occurs in the absence of α-syn aggregation within substantia nigra pars compacta dopaminergic neurons or reduced DAT terminals of the dorsal striatum, providing evidence of Lewy Pathology as a pathological process in PD and not merely an epiphenomenon of dopaminergic dysfunction. Together, these data suggest that presynaptic α-syn aggregates are sufficient to cause physiological impairments at early timepoints after initiation of aggregation.

M2-PFF induced p-α-syn aggregates in the striatum strongly colocalized to vGLUT1-positive presynaptic terminals, with minimal overlap with DAT-positive presynaptic terminals. These results support prior research that the majority of Lewy pathology in the striatum is neuritic.^14^ We previously showed using PFF injections into the striatum that intrinsic SPN excitability was decreased.^15^ In this study, there were no changes to neuron intrinsic excitability. Because α-syn aggregates localize to SPN soma and corticostriatal terminals after intrastriatal PFF injection, but not after M2-PFF injections, we conclude that the somatodendritic localization of α-syn aggregates within SPNs contributed to reduced excitability in our previous work, and suggests that pre-synaptic changes in synaptic efficacy do not drive post-synaptic changes in intrinsic properties of neurons. Both studies however, using optogenetics or electrode stimulation, showed significantly reduced evoked ESPCs and decreased asynchronous EPSC frequency, suggesting these phenotypes result from excitatory presynaptic aggregates.

Corticostriatal function is known to be critical for learning, decision making, and the consolidation of learned tasks into habits.^26,32^ The only cortical brain area to show degeneration in PD is the preSMA, suggesting selective vulnerability.^6^ Although M2 showed no degeneration six weeks after PFF injections, the selective vulnerability of this cortical area in PD makes it an important area to study, and vulnerability to α-syn aggregates compared to other cortical areas could be explored in future studies. M2-striatal function is necessary for planning and completion of sequential motor tasks.^25^ In primates, the preSMA, homologous to the M2 cortex in mice, is active during both movements of a two-movement motor sequence, initially in a reactive manner, but changing to predictive encoding over time.^9^ Studies employing these sequential motor tasks in PD patients demonstrate slowness of movement that exceeds the bradykinesia seen in single movement tasks, even when subjects were on dopamine replacement therapy during the study.^33^ Our results support the hypothesis that this sequence effect is at least partially independent of dopamine input and may rather be due to α-syn induced corticostriatal synaptic dysfunction which can be explored in the future.

The loss of glutamatergic drive in corticostriatal projections observed in the present study is concordant with our prior work and with the current literature.^15,34^ While the role of dopamine denervation cannot be ruled out in these prior studies, our data in the current study demonstrate that this glutamatergic dysfunction is independent of dopamine terminal loss. However, the mechanism by which corticostriatal synapses are impaired is still unknown. The loss of oeEPSCs observed was robust and may not be entirely accounted for by the statistically significant, yet small effect size, of vGLUT1/Homer1+ synaptic loss seen in M2-PFF injected mice. Thus, our electrophysiological data indicate that synaptic function may be impaired in M2-PFF injected mice even prior to synaptic degeneration. One potential explanation for this loss of corticostriatal projection excitability is the role of α-syn in soluble N-ethylmaleimide-sensitive factor attachment protein receptor (SNARE) complex activity. SNARE proteins are necessary for the fusion of vesicles to the plasma membrane so that neurotransmitters may be released from the presynaptic terminal. Endogenous α-syn is known to bind to vesicular SNARE VAMP2 and chaperone the complex to facilitate neurotransmitter release.^35^ Thus, aggregated α-syn may reduce the amount of endogenous α-syn available to properly facilitate neurotransmitter release.^36,37^ Here, even at early timepoints, recruitment of endogenous α-syn into phosphorylated aggregates may reduce release frequency at glutamatergic presynaptic terminals, as evidenced by the increased inter-event interval of aEPSCs in the dorsal medial striatum of M2-PFF injected mice.

The role of Lewy pathology formation in PD and DLB pathogenesis is still being uncovered, but the use of progressive models of α-syn aggregation such as the PFF model reveal that early corruption of endogenous α-syn causes neuronal dysfunction. Here, we demonstrate pathological α-syn is sufficient to drive early defects in corticostriatal synaptic transmission and loss of vGLUT1+ synaptic volume density. These findings improve our understanding of cortical defects in these synucleinopathies that may underly cognitive and other symptoms that are not ameliorated by dopamine replacement therapies. While further work is needed to determine specifically how this corticostriatal dysfunction relates to PD and DLB symptomology, targeting α-syn aggregation-mediated excitatory synaptic dysfunction could provide novel avenues for therapeutic development in LBDs.

## Methods and Materials

All details of materials used are listed in a Key Resources Table on Zenodo: **<u>10.5281/zenodo.20292044</u>**. Protocols associated with this study can be found: https://dx.doi.org/10.17504/protocols.io.ewov1rpzolr2/v1

### Mice

C57BL/6J wildtype mice (strain 000664, RRID:IMSR_JAX:000664) were obtained from the Jackson Laboratory. Mice were maintained under a 12-hour light/dark cycle and provided ad libitum access to food and water, following the National Institute of Health (NIH) guidelines for the care and use of research animals. All experimental procedures were approved by the respective Institutional Animal Care and Use Committees (IACUC) of the University of Alabama at Birmingham and the University of Florida. Male mice were used for the current study.

Unless otherwise mentioned, all reagents were purchased from Fisher Scientific.

### Preparation of **α**-syn preformed fibrils (PFF)

Mouse monomeric α-syn was expressed and purified in *E. coli* as previously described.^15^ Endotoxin was removed using Pierce High-Capacity Endotoxin Removal Spin Column, and residual endotoxin levels were measured with the Limulus amebocyte lysate (LAL) endotoxin assay kit, with levels at 0.08 to 0.13 EU/µg protein. Protein concentration was determined through 280 nm absorbance using an extinction coefficient (L) 7450 M^−1^cm^−1^ for mouse monomeric α-syn. To produce preformed fibrils (PFFs), monomeric α-syn (300 μm) was incubated at 37 °C with constant shaking for seven days in a buffer containing 150 mM potassium chloride and 50 mM Tris-HCl (pH 7.5). PFFs were separated from soluble protein by centrifugation at 21,100 x *g* for 10 minutes at room temperature (RT), then resuspended and diluted to 300 μm in the same buffer. Fragmentation into smaller, seeding-competent fibrils was performed by sonication using a Qsonica 700 W cup horn sonicator for 15 minutes at 30% amplitude with a three-second on, two-second off pulse cycle using a water chiller to maintain temperature at 15°C. The resulting fibril fragments, sized between 20 nm and 80 nm, were verified by dynamic light scattering (Dynapro Nanostar, Wyatt Technologies) as part of the quality control routine. Fragmented PFFs were subsequently used for experiments, all of which adhered to strict safety and decontamination protocols described in Fielding et al.^38^

### Stereotaxic surgeries

Intracranial injections were conducted on 3–4-month-old C57BL/6J mice using a digital stereotaxic frame. Mice received 0.1 mg/kg buprenorphine subcutaneously before the start of surgery and anesthesia was maintained with 1-3% inhaled isoflurane during the surgery; respiration was continuously monitored throughout the surgery. A gas-tight 26s-gauge point-style Hamilton syringe, controlled by a digital pump, was used to deliver 2 µL PFFs (5µg/µL), monomeric α-syn (5µg/µL) as a control, or phosphate-buffered saline (PBS) vehicle solution to the right hemisphere. Coordinates for M2 cortex were: ML: +1.5; AP: +2.5 from bregma and DV: −1.3 from dura. Injections were performed at a rate of 0.5 μL/min, and the needle was held in place for five minutes post-injection before being slowly withdrawn. Twenty-four hours after surgery, mice were administered 5mg/kg carprofen subcutaneously.

For electrophysiological recordings, mice were co-injected with 10 μg of 5 mg/mL PFFs or monomer in addition to 0.5 μL of AAV9-hSyn-ChrimsonR-tdTOMATO (Addgene), encoding the light gated ion channel ChrimsonR.

### Immunofluorescence

#### Transcardial perfusions

Mice were deeply anesthetized using isoflurane and transcardially perfused with ice-cold 0.9% saline containing 0.5% (m/v) sodium nitroprusside and 10 units/mL heparin, followed by perfusion with 4% paraformaldehyde (PFA) freshly prepared in PBS. Brains underwent post fixation in 4% PFA at 4°C overnight. They were then immersed in 30% sucrose in PBS at 4°C until the brains dropped in the solution (1-2 days), snap frozen, and stored at –80 °C. Tissue was sectioned at a thickness of 40 µm using a freezing microtome (Leica SM2010R) and preserved in cryoprotectant solution in tris-buffered saline (TBS; 50% glycerol v/v, 0.01% sodium azide w/v) at −20°C.

#### Immunofluorescence of Brain Sections

All antibodies used for immunofluorescence are listed in the Resource table. Tissue sections were rinsed three times in TBS and subjected to antigen retrieval using a solution of 10 mM sodium citrate with 0.05% Tween-20 (pH 6.0) for one hour at 37 °C. Blocking and permeabilization were performed at 4°C for one hour using a solution containing 5% goat serum and 0.1% Triton X-100 in TBS. Primary and secondary antibodies were prepared in 5% goat serum in TBS. Sections were incubated overnight at 4°C in primary antibody solution with gentle shaking. Following three washes in TBS, sections were treated with Alexa-Fluor-conjugated secondary antibodies for two hours at 4°C. Finally, sections were mounted onto Superforst Plus glass slides using Prolong Gold mounting media. Post-hoc sections from electrophysiological recordings were processed as described above but not mounted on glass slides due to tissue thickness, and instead immediately imaged using 35 mm glass-bottom MatTek dishes.

#### Immunohistochemistry of Brain Sections

Tissue sections were rinsed three times in TBS followed by immersing sections in 3% H_2_O_2_ solution for 10 min at RT. After three rinses with TBS, sections underwent antigen retrieval procedure in 10 mM sodium citrate solution for one hour at 37 °C. Sections were then rinsed in TBS and placed in blocking solution (5% donkey serum, 0.1% TritonX-100, in TBS) for one hour at 4 °C. Sections were then incubated in SatB2 (Resource table) primary antibody solution (5% donkey serum in TBS) overnight at 4 °C. After 3X TBS, sections were incubated in secondary antibody solution (5% donkey serum, 0.1% TritonX-100, in TBS) containing a biotinylated secondary antibody for two hours at 4 °C. Sections were rinsed and incubated in avidin-biotin complex (ABC) solution (ABC peroxidase kit) for one hour at RT to facilitate the formation of the avidin-biotin-complex. Sections were then developed in DAB chromogen solution for 2-3 minutes per slice, rinsed in TBS, and mounted onto glass slides (Superfrost Plus) and allowed to dry overnight at RT. Mounted samples underwent dehydration procedure in the following solutions and sequence: 30 sec H_2_O, three min 70% EtOH, three min 95% EtOH, three min 95% EtOH, three min 100% EtOH, three min 100% EtOH, five min histoclear solution, five min histoclear solution, five min histoclear solution. Dehydrated samples were coverslipped using permount mounting media and allowed to dry for at least three days at RT before microscopy.

### Microscopy and Image analysis

#### Brightfield Imaging

Whole slide tiled images to visualize SatB2 staining were generated using a Cicero Spinning disk microscope with the Nikon Ti2 base set-up (Nikon Instruments Inc). Single images were obtained in brightfield mode with a 4X air objective (Nikon CFI Plan Fluor 4X/0.13) and stitched in NiS advanced software (NiS AR V6.10.01, Nikon Instruments Inc., RRID:SCR_014329) using the large image function.

#### Stereology

Unbiased stereological analysis of SatB2-positive neurons in layer V and layer VI in M2 was conducted on an upright widefield microscope with brightfield function (Olympus, BX51) using an optical fractionator probe (Version 2023 Stereo investigator software, MBF Biosciences LLC, RRID:SCR_002526). A total of six slices (from AP+2.5mm to AP+1.0mm) with 240 μm spacing between samples were used to cover M2 cortex. To delineate the secondary motor cortices, samples were visualized using a 4X air objective (Olympus UPlan FL N 4X/0.13), and boundaries were drawn around the deep layers of M2 cortex for ipsilateral and contralateral sides. Neurons were live counted at high magnification (40X air objective, Olympus UPlan FL N 40X/0.75) using a 40X40 μm counting frame area with 22 μm optical dissector height and 200X200 μm distance between counting frames.^22^ Counting variability was assessed for each hemisphere and mouse, and Gundersen coefficient of error (CE below 0.12) was used to assess variability.

#### Confocal Microscopy

An inverted Nikon Ti2 Eclipse microscope base (Nikon Instruments Inc.) equipped with a Cicero spinning disc confocal setup (CrestOptics) was used for all confocal imaging.

##### Whole hemisphere imaging

For visualization of p-α-syn-positive aggregates throughout whole coronal sections, large tiled (10% overlap between tiles) images were acquired with a 20X air objective (Nikon CFI Plan Fluor 20X/0.5). Brightness and contrast were adjusted using NiS-Elements Advanced Research software (NiS AR V6.10.01, Nikon Instruments Inc., RRID:SCR_014329) by adjusting look-up table (LUT) settings and exporting images as tiff files.

##### Dopamine Transporter (DAT) fiber density

DAT and NeuN fluorescent signal were acquired in the dorsal and ventral striatum for ipsilateral and contralateral hemispheres using a 20X air objective (Nikon CFI Plan Fluor 20X/0.5). Five single plane images per region of interest (dorsal or ventral, ipsilateral or contralateral) were acquired in five separate coronal sections covering the striatum. Images were acquired using full sensor size with no frame averaging, while keeping light source intensities and exposure times per channel constant throughout the experiment.

##### Synapse analysis

Brain sections were imaged using a 60X oil immersion objective (Nikon CFI Plan Apochromat λD 60X/1.42). Ten z-stacks were collected for at least two separate coronal slices in the ipsilateral dorsal striatum (AP 0.9-0.00mm from bregma) of each mouse (2 µm z-stacks, step size 0.125 µm) using full sensor size (2048X2048) without frame averaging. Light source intensities and exposure times per channel were kept constant throughout the experiment. To prevent surface stain artifacts, z-stacks were acquired at 4 µm depth from surface.

##### Colocalization analysis

To prevent false positive results for intensity-based colocalization analysis, only single plane images were acquired. Ten single plane images of the dorsal striatum were acquired per mouse using a 60X oil immersion objective (Nikon CFI Plan Apochromat λD 60X/1.42) at full sensor size. Light source intensities and exposure times per channel were kept constant throughout the experiment.

#### Image Analysis

##### Synapse analysis

Synapse analyses were performed as described.^24^ Confocal z-stacks were deconvolved using Lucy-Richardson algorithm (20 iterations) batch deconvolution using NiS-Elements Advanced Research software (NiS AR V6.10.01, Nikon Instruments Inc., RRID:SCR_014329). Deconvolved z-stacks were converted into IMS files using Imaris software (V.10.2, Bitplane via Oxford Industries, RRID:SCR_007370), and surfaces were detected. Using a shortest distance filter, synaptic loci surfaces were defined as pre and postsynaptic surfaces with shortest distance below 0.01 µm. Total and synaptic surface densities were calculated by dividing surface count by total z-stack volume. To compute total z-stack volumes, a surface filling the entire z-stack was generated and its volume measured. For morphology parameters, mean values for each z-stack were averaged, and all mean values averaged per mouse. All Imaris surface construction and analysis parameters were kept constant within experiments.

##### DAT fiber density

Fluorescent intensity of DAT and NeuN were measured in Fiji/Image J (RRID: RRID:SCR_002285), and DAT intensity was normalized to NeuN intensity per image frame. Mean values per mouse were computed from five independent imaging frames.

##### Colocalization analysis

Colocalization between presynaptic markers and p-α-syn-positive aggregates was assessed via thresholded Mander’s Colocalization Coefficient (tM1, p-α-syn over vGLUT1, or p-α-syn over DAT) using the coloc2 plugin for Fiji/Image J (RRID:SCR_003070). Ten images per mouse were analyzed and mean values calculated.

### *Ex vivo* slice recordings

#### Preparation of slices

Unless otherwise stated, reagents were purchased from Sigma Aldrich. After deep anesthesia with isoflurane, mice were perfused with cold 95% O_2_ and 5% CO_2_ gas-equilibrated N-Methyl-D-gluconate solution (93 mM NMDG, 2.5 mM KCl, 1.4 mM Na-Phosphate Monobasic Monohydrate, 30 mM Na-bicarbonate, 20 mM Hepes, 5 mM Glucose, 2 mM thiourea, 3 mM Na-Pyruvate, 10 mM MgSO_4_, 0.5 mM CaCl_2_) as described in Chambers et al. 2024 and previous work.^15,39^ Coronal slices (250 µm thick) containing the striatum were sectioned on a vibratome (VT1200S, Leica) while maintaining immersion in cold, gas-equilibrated NMDG solution. Sections were recovered in warm (33 °C), gas equilibrated NMDG solution for 10 minutes, followed by one hour recovery at RT in a sodium ascorbate (5 mM) and gas-equilibrated artificial cerebrospinal fluid (aCSF) solution (126 mM NaCl, 2.5 mM KCl, 1.4 mM Na-Phosphate Monobasic Monohydrate, 26 mM Na-bicarbonate, 1 mM Glucose, 1.5 mM MgSO_4_, 2 mM CaCl_2_, 300-205 mOsm) until use.

#### Electrophysiological recordings

Coronal sections in the recording chamber were perfused with warm, gas-equilibrated aCSF at a rate of 2 mL/min while maintaining a solution temperature of 33 °C using an in-line heating element. IR-DIC Optics with a DAGE-2000 infrared camera was used to visualize neurons. To identify neurons receiving innervation from ChrimsonR-positive M2-terminals, tdTOMATO-positive terminals were visualized using a combination of green LED light and red-light filter cubes settings allowing for visualization of fluorescent signal. Glass borosilicate pipettes (3-5 MΩ resistance) filled with internal K-gluconate solution (125 mM K-gluconate, NaCl 4mM, 10 mM HEPES, 4 mM Mg-ATP, 0.3 mM Na-ATP, 10 mM Tris-phosphate) were used for whole cell patch clamp recordings of SPNs. Electrophysiological data was acquired using a Multiclamp 700B amplifier (Molecular Devices) and digitized at 10 kHz using Axon Digidata 1550B (Molecular Devices) with pClamp software (V11.2 Molecular Devices, RRID:SCR_011323). The acquired data was then filtered using a low pass filter at 1 kHz and a high pass filter at 10 kHz. SPNs were identified by meeting the following criteria: Resting membrane potential around –80 mV, no spontaneous action potential (AP) firing, and no frequency adaptation of AP firing upon depolarization. If access resistance exceeded 30 MΩ, recordings were rejected from analysis. To assess spontaneous excitatory postsynaptic currents (sEPSC), SPNs were recorded for five minutes in voltage clamp configuration (holding potential: −75 mV). Next, SPNs were depolarized with current injections (50 pA increment, from 0 to 700 pA, 1 s depolarization, 1 s recovery) in current clamp. M2-to-striatum optogenetically evoked glutamate transmission experiments were performed using far-red LED light stimulation of ChrimsonR-positive M2 terminals and measuring postsynaptic optogenetically-evoked (oe) EPSC in SPNs held at –75 mV. LED light stimuli at 0.5, 1.0, 3.3 and 8.0 mW/mm^2^ (irradiance was empirically determined via a light meter) were applied as a paired-pulse (50 ms interval). Paired pulse ratio (PPR) was calculated by peak amplitude from Stimulus 2 divided by the peak amplitudes for Stimulus 1. For each cell recorded at each stimulus the PPR was applied five times. For asynchronous release experiments, slices were perfused with aCSF solution containing 2 mM SrCl_2_, substituting CaCl_2_. After the recordings concluded, sections were immersed in 4% PFA in PBS for 30 minutes at RT to allow for post fixation of the tissue. Post hoc staining of fixed electrophysiology slices was performed as described in *Immunofluorescence of Brain Sections*.

#### Electrophysiology data analysis

All electrophysiological data analysis was performed in Clampfit software (part of pClamp software suite, V11.2 Molecular Devices, RRID:SCR_011323).

##### sEPSC

Spontaneous excitatory currents of five-minute recordings were performed using the detect event function via template search in Clampfit Software. The same sEPSC template was used in all recordings for semi-automatic event detection. To assess spontaneous transmission, inter event interval (IEI) for overall activity and EPSC amplitudes were measured.

##### OeEPSC and PPR

After baseline subtraction, EPSC amplitudes were measured using the event detection function for first and second stimulation. The amplitude of the second EPSC peak was divided by amplitude of first EPSC peak to assess the PPR.

##### aEPSC

As described for sEPSC, IEI and EPSC amplitude were analyzed by using Clampfit software event detection tool.

### Statistics

Graph Pad Prism software (V.10.0.2, RRID:SCR_002798) was used for all statistical analysis and generation of graphical data illustration. All data is represented as mean±SEM if not otherwise stated in figure legends. Data were explored for normality using D’Agostino & Pearson or Shapiro-Wilk tests. Homoscedasticity of variance between groups was evaluated with the Brown-Forsythe test, and if data sets violated equality of variance, Brown-Forsythe or Welch correction were applied. Welch correction was applied for datasets with unequal variation.

#### Unpaired Student t-test

If data fit a normal distribution, two-tailed unpaired t-tests were used to assess statistical differences (alpha set to 0.05) between two groups. Analyses of non-normally distributed data were performed by Mann-Whitney testing.

#### One-way ANOVA

If the data fit a normal distribution, one-way ANOVA test was applied with Tukey’s multiple comparisons test. Kruskal-Wallis test was used for analysis of non-normally distributed datasets.

#### Two-way ANOVA

For grouped analysis with two independent factors or more, two-way ANOVA was used to assess for significant row or column effects, or significant interactions.

#### Repeated measure ANOVA

Data from experiments with repeated measure design were analyzed with either two-way repeated measure ANOVA or mixed effect model.

## Data Availability

The data, protocols, and key lab materials used and generated in this study are listed in a Key Resources Table alongside their persistent identifiers at **10.5281/zenodo.20292044**.

The source data are deposited on Zenodo **10.5281/zenodo.20292044.**

All raw .nd2 image files are deposited on BioStudies 10.6019/S-BIAD2548.

No code was generated for this study; all data cleaning, preprocessing, analysis, and visualization was performed using Excel and Graphpad Prism.

## Acknowledgements

This research was funded in whole or in part by Aligning Science Across Parkinson’s [020616] through the Michael J Fox Foundation [023031] to LVD and MMohele, and Parkinson’s Foundation visiting fellow award to CB. For the purpose of open access, the author has applied a CC-BY 4.0 public copyright license to all Author Accepted Manuscripts arising from this submission.

## Author Contributions

CB collected data, analyzed data, generated figures, wrote the first draft of the manuscript and helped revise later version; ZF wrote the abstract, introduction and discussion, analyzed data, interpreted data, generated figures; MMenard collected data, organized and submitted data for open access, wrote protocols, revised the manuscript; HC, JH, DN, IG, MMillet assisted with experiments and data analysis; JAH interpreted data, wrote and revised manuscript; MMoehle designed experiments, supervised the project, analyzed data and interpreted data, revised the manuscript; LVD designed experiments, supervised the project, analyzed and interpreted data, wrote and revised the manuscript, revised figures, assisted in organizing data and writing protocols for open access submission.

The authors declare no competing interests.

## References

1 Braak, H. et al. Staging of brain pathology related to sporadic Parkinson’s disease. Neurobiol Aging 24, 197–211 (2003). 10.1016/s0197-4580(02)00065-9

2 Beach, T. G. et al. Unified staging system for Lewy body disorders: correlation with nigrostriatal degeneration, cognitive impairment and motor dysfunction. Acta Neuropathol 117, 613–634 (2009). 10.1007/s00401-009-0538-8

3 Graves, S. M. & Surmeier, D. J. Delayed Spine Pruning of Direct Pathway Spiny Projection Neurons in a Mouse Model of Parkinson’s Disease. Front Cell Neurosci 13, 32 (2019). 10.3389/fncel.2019.00032

4 Villalba, R. M. & Smith, Y. Loss and remodeling of striatal dendritic spines in Parkinson’s disease: from homeostasis to maladaptive plasticity? J Neural Transm (Vienna) 125, 431–447 (2018). 10.1007/s00702-017-1735-6

5 Haslinger, B. et al. Event-related functional magnetic resonance imaging in Parkinson’s disease before and after levodopa. Brain 124, 558–570 (2001). 10.1093/brain/124.3.558

6 MacDonald, V. & Halliday, G. M. Selective loss of pyramidal neurons in the pre-supplementary motor cortex in Parkinson’s disease. Mov Disord 17, 1166–1173 (2002). 10.1002/mds.10258

7 Rahimpour, S., Rajkumar, S. & Hallett, M. The Supplementary Motor Complex in Parkinson’s Disease. J Mov Disord 15, 21–32 (2022). 10.14802/jmd.21075

8 Hanoglu, L., Saricaoglu, M., Toprak, G., Yilmaz, N. H. & Yulug, B. Preliminary findings on the role of high-frequency (5Hz) rTMS stimulation on M1 and pre-SMA regions in Parkinson’s disease. Neurosci Lett 724, 134837 (2020). 10.1016/j.neulet.2020.134837

9 Nakajima, T., Hosaka, R. & Mushiake, H. Complementary Roles of Primate Dorsal Premotor and Pre-Supplementary Motor Areas to the Control of Motor Sequences. J Neurosci 42, 6946–6965 (2022). 10.1523/JNEUROSCI.2356-21.2022

10 Battaglia, S., Nazzi, C., Di Fazio, C. & Borgomaneri, S. The role of pre-supplementary motor cortex in action control with emotional stimuli: A repetitive transcranial magnetic stimulation study. Ann N Y Acad Sci 1536, 151–166 (2024). 10.1111/nyas.15145

11 Hendrix, C. M. et al. Parkinsonism Disrupts Neuronal Modulation in the Presupplementary Motor Area during Movement Preparation. J Neurosci 45 (2025). 10.1523/JNEUROSCI.1802-24.2025

12 Benecke, R., Rothwell, J. C., Dick, J. P., Day, B. L. & Marsden, C. D. Disturbance of sequential movements in patients with Parkinson’s disease. Brain 110 (Pt 2), 361–379 (1987). 10.1093/brain/110.2.361

13 Werheid, K., Koch, I., Reichert, K. & Brass, M. Impaired self-initiated task preparation during task switching in Parkinson’s disease. Neuropsychologia 45, 273–281 (2007). 10.1016/j.neuropsychologia.2006.07.007

14 Duda, J. E., Giasson, B. I., Mabon, M. E., Lee, V. M. & Trojanowski, J. Q. Novel antibodies to synuclein show abundant striatal pathology in Lewy body diseases. Ann Neurol 52, 205–210 (2002). 10.1002/ana.10279

15 Brzozowski, C. F. et al. Early alpha-synuclein aggregation decreases corticostriatal glutamate drive and synapse density. Neurobiology of disease 210, 106918 (2025). 10.1016/j.nbd.2025.106918

16 Tozzi, A. et al. Dopamine-dependent early synaptic and motor dysfunctions induced by alpha-synuclein in the nigrostriatal circuit. Brain 144, 3477–3491 (2021). 10.1093/brain/awab242

17 Weber, M. A. et al. Alpha-Synuclein Pre-Formed Fibrils Injected into Prefrontal Cortex Primarily Spread to Cortical and Subcortical Structures. J Parkinsons Dis 14, 81–94 (2024). 10.3233/JPD-230129

18 Bieri, G., Gitler, A. D. & Brahic, M. Internalization, axonal transport and release of fibrillar forms of alpha-synuclein. Neurobiology of disease 109, 219–225 (2018). 10.1016/j.nbd.2017.03.007

19 Volpicelli-Daley, L. A. et al. Exogenous alpha-synuclein fibrils induce Lewy body pathology leading to synaptic dysfunction and neuron death. Neuron 72, 57–71 (2011). 10.1016/j.neuron.2011.08.033

20 Delic, V. et al. Sensitivity and specificity of phospho-Ser129 alpha-synuclein monoclonal antibodies. J Comp Neurol 526, 1978–1990 (2018). 10.1002/cne.24468

21 Hintiryan, H. et al. Connectivity characterization of the mouse basolateral amygdalar complex. Nat Commun 12, 2859 (2021). 10.1038/s41467-021-22915-5

22 Stoyka, L. E. et al. Behavioral defects associated with amygdala and cortical dysfunction in mice with seeded α-synuclein inclusions. Neurobiology of disease 134, 104708 (2020). 10.1016/j.nbd.2019.104708

23 Goralski, T. M. et al. Spatial transcriptomics reveals molecular dysfunction associated with cortical Lewy pathology. Nat Commun 15, 2642 (2024). 10.1038/s41467-024-47027-8

24 Gcwensa, N. Z., Long, K. Y., Manabat, A. F. & Volpicelli-Daley, L. A. Protocol to study synapse density or volume-SynDOVE-in brain using confocal microscopy and Imaris three-dimensional surface rendering software. STAR Protoc 7, 104465 (2026). 10.1016/j.xpro.2026.104465

25 Rothwell, P. E. et al. Input- and Output-Specific Regulation of Serial Order Performance by Corticostriatal Circuits. Neuron 88, 345–356 (2015). 10.1016/j.neuron.2015.09.035

26 Pennartz, C. M. B., Joshua D; Graybiel, Ann M; Ito, Rutsuko; Lansink, Carien S; van der Meer, Matthijs; Redish, A David; Smith, Kyle S; Voorn, Pieter. Corticostriatal Interactions during Learning, Memory Processing, and Decision Making. J Neurosci 29, 12831–12838 (2009). 10.1523/JNEUROSCI.3177-09.2009

27 Day, M. et al. Selective elimination of glutamatergic synapses on striatopallidal neurons in Parkinson disease models. Nat Neurosci 9, 251–259 (2006). 10.1038/nn1632

28 Paille, V. et al. Distinct levels of dopamine denervation differentially alter striatal synaptic plasticity and NMDA receptor subunit composition. J Neurosci 30, 14182–14193 (2010). 10.1523/JNEUROSCI.2149-10.2010

29 Froula, J. M. et al. Defining alpha-synuclein species responsible for Parkinson’s disease phenotypes in mice. The Journal of biological chemistry 294, 10392–10406 (2019). 10.1074/jbc.RA119.007743

30 Gcwensa, N. Z. et al. Excitatory synaptic structural abnormalities produced by templated aggregation of alpha-syn in the basolateral amygdala. Neurobiology of disease 199, 106595 (2024). 10.1016/j.nbd.2024.106595

31 Sah, S. et al. Progressive vulnerability of cortical synapses in alpha-synucleinopathy. bioRxiv (2025). 10.1101/2024.06.20.599774

32 Graybiel, A. M. Habits, rituals, and the evaluative brain. Annual Review of Neuroscience 31, 359–387 (2008). 10.1146/annurev.neuro.29.051605.112851

33 Avanzino, L., Pelosin, E., Martino, D. & Abbruzzese, G. Motor timing deficits in sequential movements in Parkinson disease are related to action planning: a motor imagery study. PLoS One 8, e75454 (2013). 10.1371/journal.pone.0075454

34 Chen, L. et al. Synaptic location is a determinant of the detrimental effects of alpha-synuclein pathology to glutamatergic transmission in the basolateral amygdala. Elife 11 (2022). 10.7554/eLife.78055

35 Wang, L. et al. alpha-synuclein multimers cluster synaptic vesicles and attenuate recycling. Curr Biol 24, 2319–2326 (2014). 10.1016/j.cub.2014.08.027

36 Sun, J. et al. Functional cooperation of alpha-synuclein and VAMP2 in synaptic vesicle recycling. Proc Natl Acad Sci U S A 116, 11113–11115 (2019). 10.1073/pnas.1903049116

37 Yoo, G., Shin, Y. K. & Lee, N. K. The Role of alpha-Synuclein in SNARE-mediated Synaptic Vesicle Fusion. J Mol Biol 435, 167775 (2023). 10.1016/j.jmb.2022.167775

38 Fielding, L. et al. Current safety recommendations for handling mouse and human alphasynuclein pre-formed fibrils. Neurobiology of disease 206, 106820 (2025). 10.1016/j.nbd.2025.106820

39 Chambers, N. E. et al. Conditional Knockout of Striatal Gnal Produces Dystonia-like Motor Phenotypes. bioRxiv (2024). 10.1101/2024.08.26.609754

