## Supplemental figures for "α-Synuclein aggregates in corticostriatal terminals impair glutamatergic transmission in the absence of neurodegeneration"

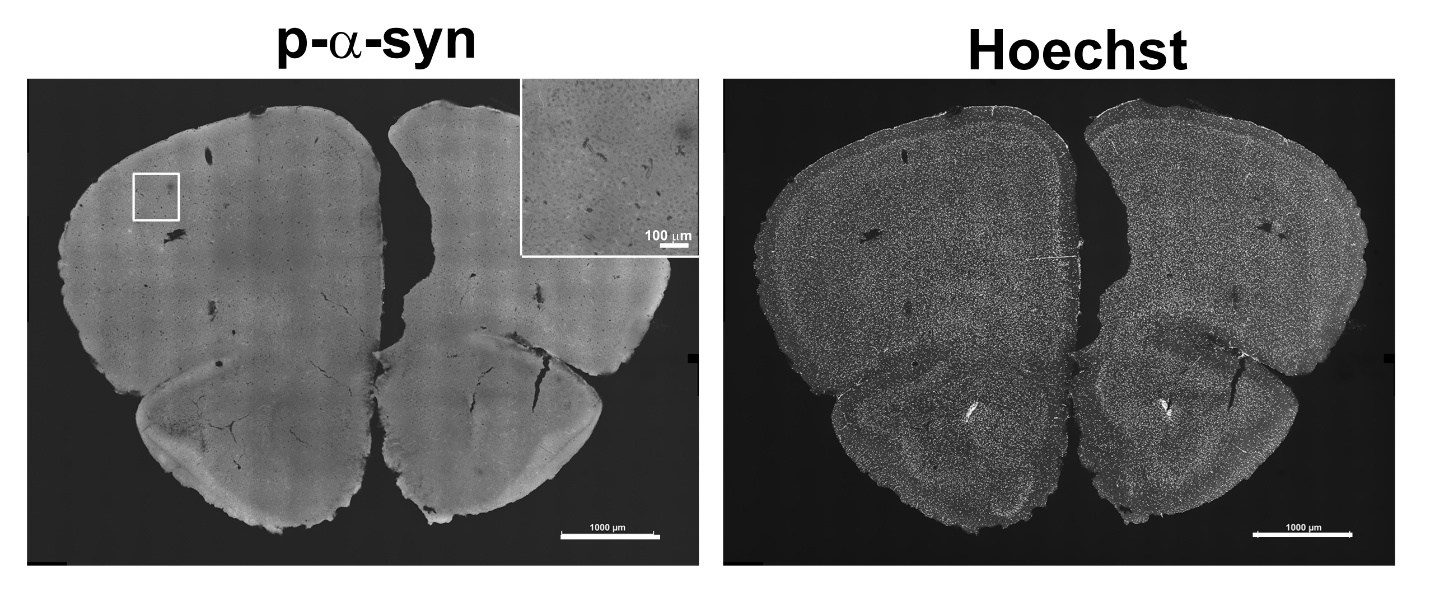

**Supplemental Figure 1.** Mice were injected with 10 µg α-syn monomer in M2 cortex and perfused six weeks later. Immunofluorescence was performed with an antibody to p-α-Syn with Hoechst as a counterstain. The coronal image is representative of rostral cortex with M2. Scale bars = 1000 µm, 100 µm.

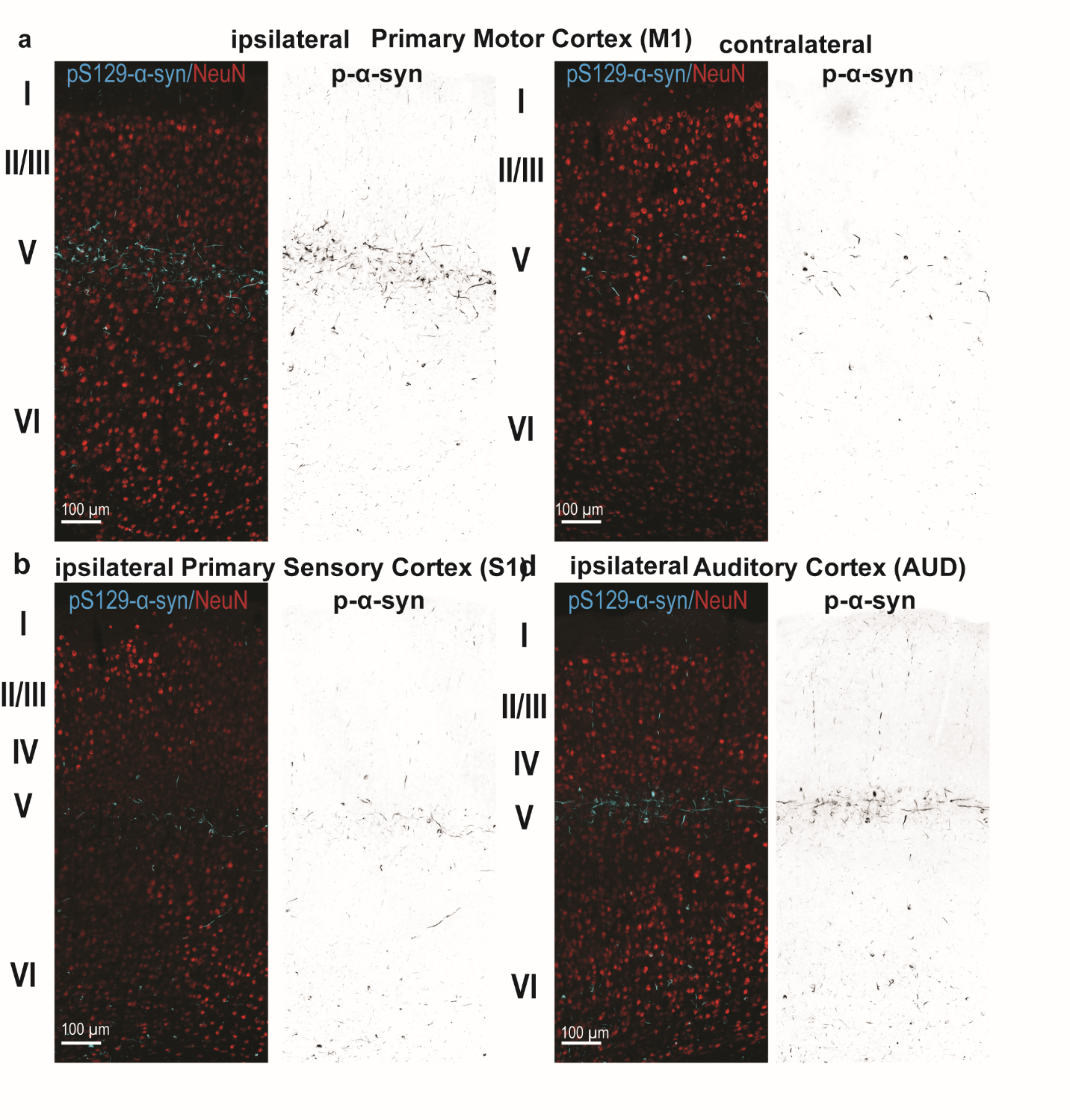

**Supplemental Figure 2.** Mice were injected with 10 µg α-syn PFFs in M2 cortex and perfused six weeks later. Immunofluorescence was performed with an antibody to p-α-syn (blue) and the neuronal marker, NeuN (red). Representative areas of the cortex with layers indicated are shown” **a)** M1 primary motor cortex ipsilateral and contralateral, **b)** ipsilateral primary sensory cortex **c)** ipsilateral auditory cortex. Inverted black and white images are shown next to the immunofluorescent images to help with visualization of p-α-syn aggregates. Scale bars = 100 µm.

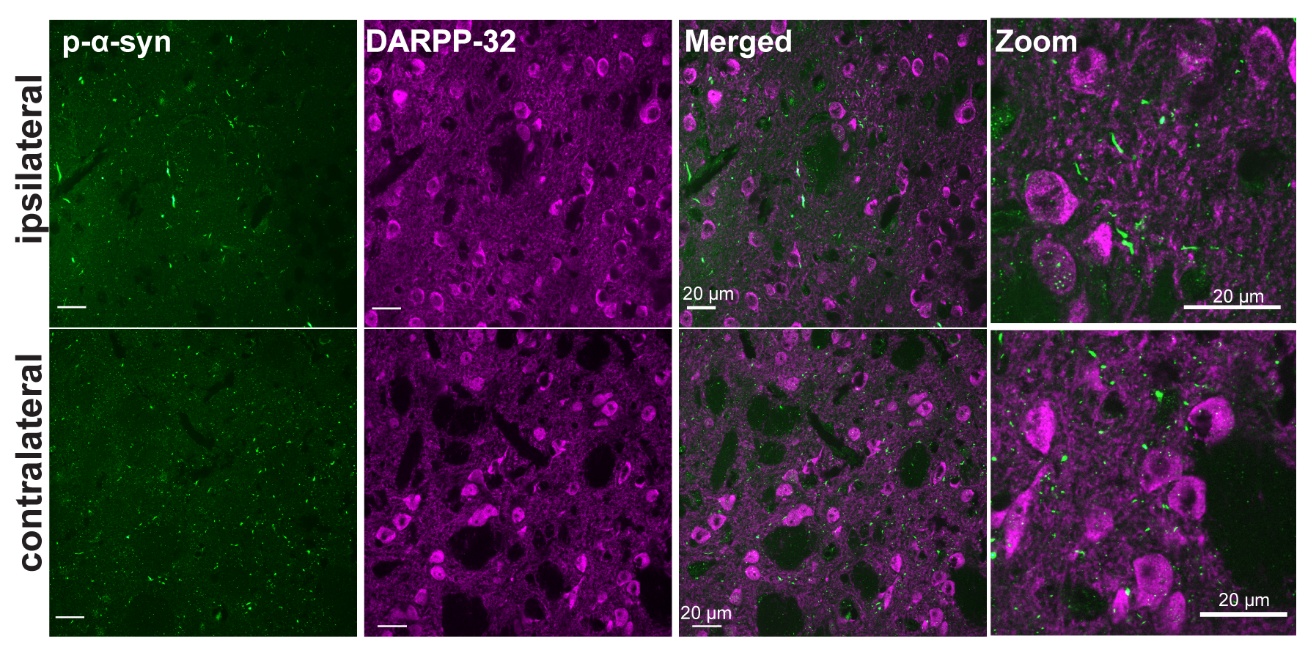

**Supplemental Figure 3.** Mice were injected with 10 µg α-syn PFFs in M2 cortex and perfused six weeks later. Immunofluorescence was performed with an antibody to p-α-syn (green) and the SPN marker, DARPP-32 (magenta). Representative images from the striatum ipsilateral and contralateral to the injection site are shown. The images in column 4 are zoomed in images from column 3. Scale bars = 20, 100 µm.

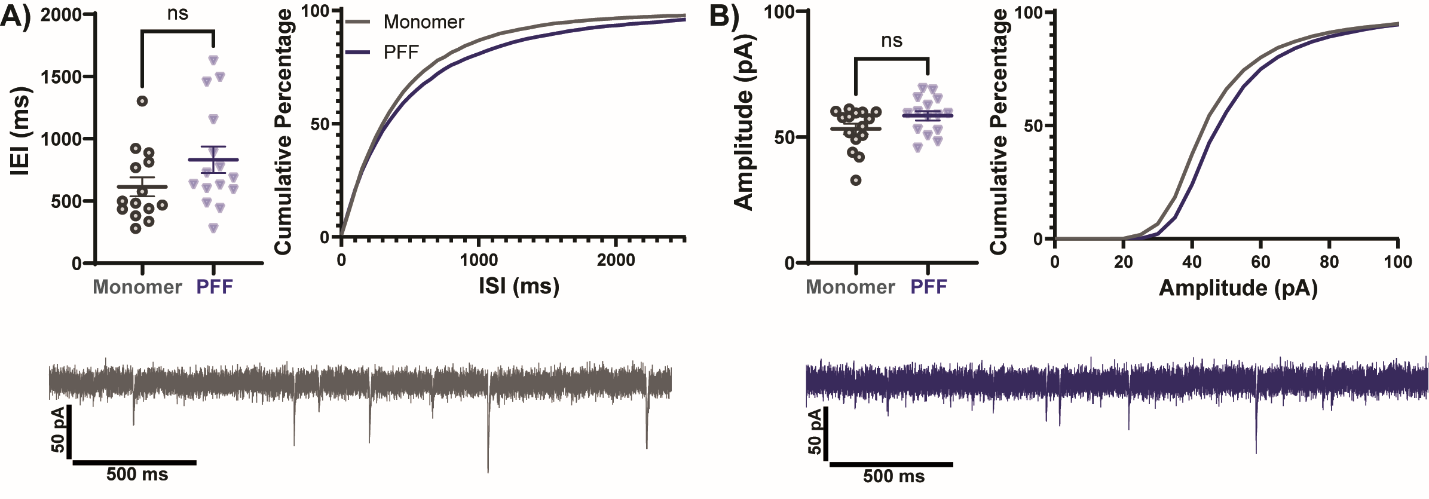

**Supplemental Figure 4.** Mice were injected with 10 µg α-syn PFFs or monomer in M2 cortex. Six weeks later whole-cell patch clamp recording of SPNs were performed in coronal slices containing striatum. A) Quantitation of median inter event interval (IEI) and cumulative probabilities are shown. No differences were observed between groups (Unpaired Student’s t-test: p= 0.8993) Monomer: N=4, PFF N=4. B) Quantitation of mean amplitude of sEPSCs and cumulative probabilities are shown. (Mann-Whitney test: p=0.137).

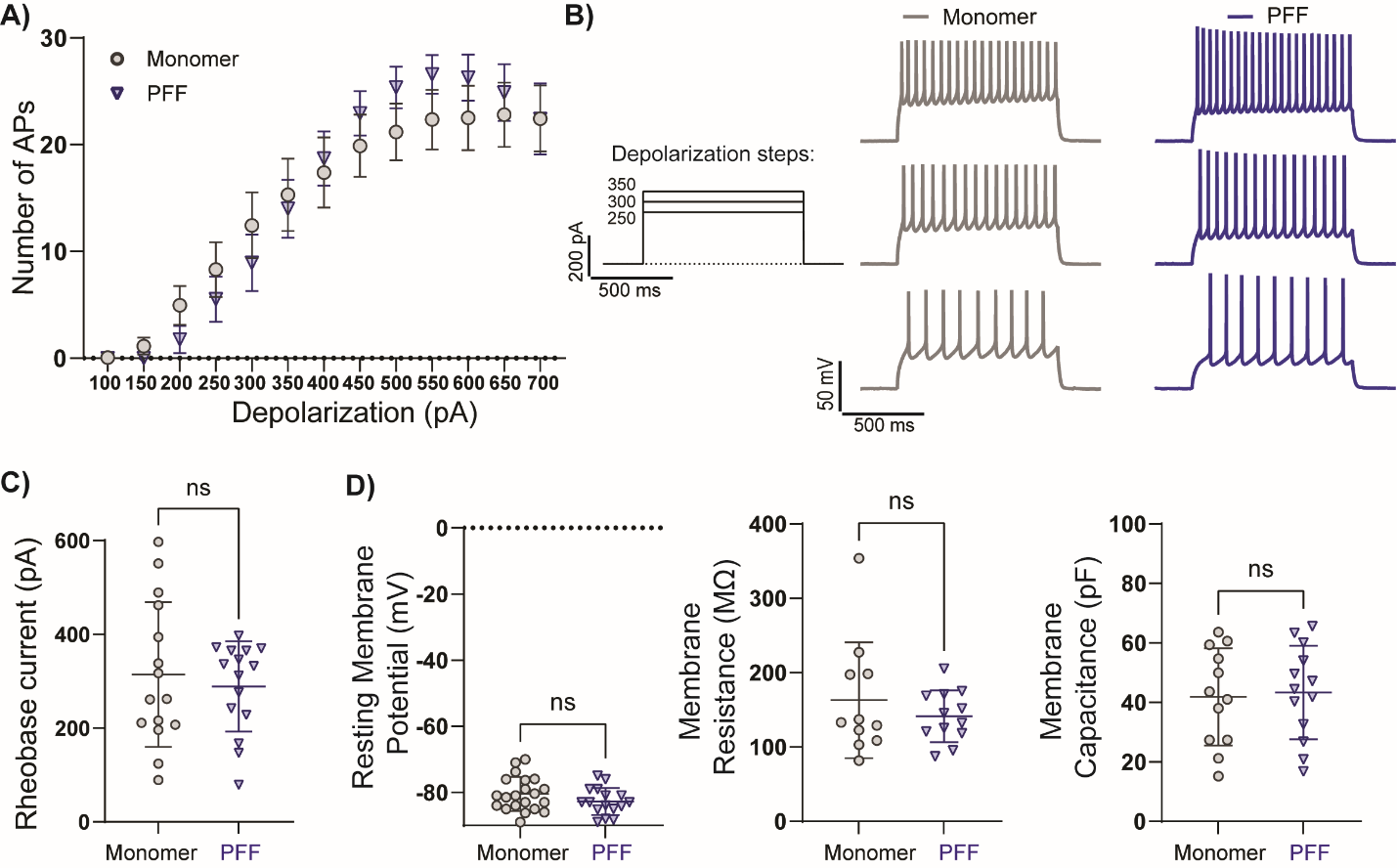

**Supplemental Figure 5.** Mice were injected with 10 µg α-syn PFFs or monomer in M2 cortex. Six weeks later whole cell patch clamp recordings of SPNs were performed in coronal slices. **a)** Numbers of action potentials elicited by 1ms depolarizing current injections. **b)** Representative voltage traces and action potentials in monomer and PFF injected mice with 250, 300, 350 pA depolarization steps (Two way ANOVA row factor (monomer vs. PFF) F(1,35)=0.06593, p=0.7989. **c)** Quantitation of rheobase current in SPNs to measure intrinsic excitability (Mann-Whitney test p=0.9589). **d)** Measures of resting membrane potentials (Unpaired t-test, p=0.159), membrane resistance (Mann-Whitney test p=0.7859), and membrane capacitance (Unpaired t test, p=0.8141).
